# Estimation of High-Order Aberrations and Anisotropic Magnification from Cryo-EM Datasets in RELION-3.1

**DOI:** 10.1101/798066

**Authors:** Jasenko Zivanov, Takanori Nakane, Sjors H.W. Scheres

## Abstract

We present methods that detect three types of aberrations in single-particle cryo-EM data sets: symmetrical and antisymmetrical optical aberrations and magnification anisotropy. Because our methods only depend on the availability of a preliminary 3D reconstruction from the data, they can be used to correct for these aberrations for any given cryo-EM data set, *a posteriori*. Using five publicly available data sets, we show that considering these aberrations improves the resolution of the 3D reconstruction when the effects are present. The methods are implemented in version 3.1 of our open-source software package RELION.

## 1. Introduction

Structure determination of biological macromolecules using electron cryo-microscopy (cryo-EM) is primarily limited by the radiation dose to which samples can be exposed before they are destroyed. As a consequence of the low electron dose, cryo-EM has to rely on very noisy images. In recent years, advances in electron-detector technology and processing algorithms have enabled the reconstruction of molecular structures at resolutions sufficient for *de novo* atomic modelling (Fernandez-Leiro & Scheres, 2016). With increasing resolutions, limitations imposed by the optical system of the microscope are becoming more important. In this paper, we propose methods to estimate three optical effects – symmetrical and antisymmetrical aberrations and magnification anisotropy – which, when considered during reconstruction, increase the attainable resolution.

In order to increase contrast, cryo-EM images are typically collected out of focus, which introduces a phase shift between the scattered and unscattered components of the electron beam. This phase shift varies with spatial frequency and gives rise to the contrast-transfer function (CTF). Since the electron-scattering potential of the sample corresponds to a real-valued function, its Fourier-space representation exhibits Friedel symmetry, i.e. the amplitude of the complex structure factor at spatial frequency *k* is the complex-conjugate of the structure factor at frequency −*k*. In an ideal microscope, the phase shift of those two frequencies would be identical, and the CTF could be expressed as a real-valued function. In reality, however, imperfections of the optical system can produce asymmetrical phase-shifts that break the Friedel symmetry of the scattered wave. The effect of this is that the CTF becomes a complex-valued function, which not only affects the amplitudes of the structure factors, but also their phases.

The phase-shifts of a pair of corresponding spatial frequencies can be separated into a symmetrical component (i.e. their average shift) and an antisymmetrical one (i.e. their deviation from that average). In this paper, we will refer to the antisymmetrical component as *antisymmetrical aberrations*. The symmetrical component of the phase shift sometimes also deviates from the one predicted by the aberration-free CTF model (Hawkes & Kasper, 1996). The effect of this is that the CTF is not always adequately represented by a set of elliptical rings of alternating sign, but the so-called Thon rings can take on slightly different shapes. We will refer to this deviation from the traditional CTF model as *symmetrical aberrations*.

In addition to the antisymmetrical and symmetrical aberrations, the recorded image itself can be distorted by a different magnification in two perpendicular directions. This is called anisotropic magnification. Anisotropic magnification can be detected by measuring the ellipticity of the power spectra of multi-crystalline test samples (Grant & Grigorieff, 2015). Provided the microscope objective lens astigmatism is small, systematic differences between the defoci in two perpendicular directions have also been proposed as a means to detect anisotropic magnification (Zhao *et al.*, 2015).

Because the antisymmetrical and symmetrical aberrations and the anisotropic magnification produce different effects, we propose three different and independent methods to estimate them. We recently proposed a method to estimate a specific type of antisymmetrical aberration that arises from a tilted electron beam (Zivanov *et al.*, 2018). In this paper, we propose an extension of that method that allows us to estimate arbitrary antisymmetrical aberrations expressed as linear combinations of Zernike polynomials. The methods to estimate symmetrical aberrations and anisotropic magnification are novel. Similar to the method for antisymmetrical aberration correction, the method for symmetrical aberration correction also uses Zernike polynomials to model the estimated aberrations. The choice of Zernike polynomials as a basis is to some degree arbitrary, and the methods described here could be trivially altered to use any other function as a basis.

Optical aberrations in the electron microscope have been studied extensively in the material science community (Batson *et al.*, 2002; Krivanek *et al.*, 2008; Saxton, 1995; Saxton, 2000; Meyer *et al.*, 2002). However, until now, their estimation has required specific test samples of known structure and of greater radiation resistance than biological samples. The methods presented in this paper work directly on cryo-EM single-particle data sets of biological samples, making it possible to estimate the effects after the data have been collected, and without performing additional experiments on specific test samples. Using data sets that are publicly available from the EMPIAR data base (Iudin *et al.*, 2016), we illustrate that when these optical effects are present, their correction leads to reconstruction with increased resolution.

## 2. Materials and Methods

### 2.1. Observation Model

We are working on a single-particle cryo-EM data set consisting of a large number of particle images. We assume that we already have a preliminary 3D reference map of the particle up to a certain resolution, and that we know the approximate viewing parameters of all observed particles. This allows us to predict each particle image, which in turn allows us to estimate the parameters of the optical effects by comparing those predicted images to the observed ones.

Let *X*_*p*,**k**_ ∈ ℂ be the complex amplitude of the observed image of particle *p* ∈ ℕ for 2D spatial frequency **k** ∈ ℤ^2^. Without loss of generality, we can assume that the observed image is shifted so that the centre of the particle appears at the origin of the image. We can obtain the corresponding predicted image by integrating over the 3D reference along the appropriate viewing direction. According to the central-slice theorem, the corresponding complex amplitude *V*_*p*,**k**_ ∈ ℂ of the predicted particle image is given by

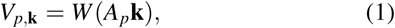

where *W* : ℝ ^3^ ↦ ℂ is the 3D reference map in Fourier space and *A*_*p*_ is a 3 × 2 projection matrix arising from the viewing angles. Since the backprojected positions of the 2D pixels **k** mostly fall between the 3D voxels of the reference map, we determine the values of *W* (*A*_*p*_**k**) using linear interpolation.

Further, we assume that we have an estimate of the defocus and astigmatism of each particle, as well as the spherical aberration of the microscope, allowing us to also predict the CTFs. We can therefore write:

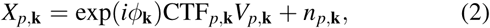

where *ϕ*_**k**_ is the phase shift angle induced by the antisymmetrical aberration, CTF_*p*,**k**_ is the real part of the CTF, and *n*_*p*,**k**_ represents the noise.

The three methods presented in the following all aim to estimate the optical effects by minimizing the squared difference between *X*_*p*,**k**_ and exp(*iϕ*_**k**_)CTF_*p*,**k**_*V*_*p*,**k**_. This is equivalent to a maximum-likelihood estimate under the assumption that all *n*_*p*,**k**_ are drawn from the same normal distribution.

### 2.2. Antisymmetrical Aberrations

Antisymmetrical aberrations shift the phases in the observed images and they are expressed by the angle *ϕ*_**k**_ in Eq. 2. We assume that *ϕ*_**k**_ is constant for a sufficiently large number of particles. This assumption is necessary since, in the presence of typically strong noise, we require the information from a large number of images to obtain a reliable estimate.

We model *ϕ*_**k**_ using antisymmetrical Zernike polynomials as a basis:

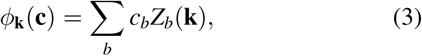

where *c*_*b*_ ∈ ℝ are the unknown coefficients describing the aberration and *Z*_*b*_(**k**) are a subset of the antisymmetrical Zernike polynomials. The usual two-index ordering of those polynomials is omitted for the sake of clarity. The coefficients *c*_*b*_ are determined by minimizing the following sum of squared differences over all particles:

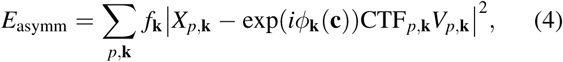

where *f*_**k**_ is a heuristical weighting term given by the FSC of the reconstruction – its purpose is to suppress the contributions of frequencies |**k**| for which the reference is less reliable.

Since typical data sets contain between 10^4^ and 10^6^ particles, and each particle image typically consists of more than 104 Fourier pixels, optimizing the non-linear expression in Eq. 4 directly would be prohibitive. Instead, we apply a two-step approach. First, we reduce the above sum over sums of quadratic functions to a single sum over quadratic functions, one for each Fourier-space pixel **k**:

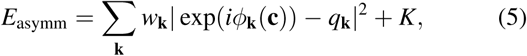

where *K* is a constant that does not influence the optimum of *c*_*b*_. The per-pixel optimal phase shifts *q*_**k**_ ∈ ℂ and weights *w*_**k**_ ∈ ℝ are given by:

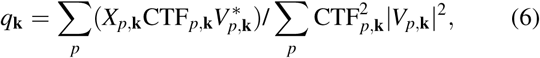

,

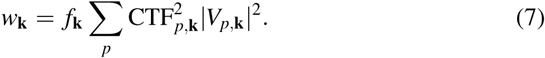

This is the same transformation that we have applied for the beam-tilt estimation in RELION 3.0 (Zivanov *et al.*, 2018) – beam tilt is in fact only one of the possible sources of antisymmetrical aberrations. The computation of *q*_**k**_ and *w*_**k**_ requires only one iteration over all the images in the data set, and it usually takes on the order of one hour of CPU time.

Once the *q*_**k**_ and *w*_**k**_ are known, the optimal *c*_*b*_ are determined by minimizing the following sum of squared differences using the Nelder-Mead downhill simplex (Nelder & Mead, 1965) method:

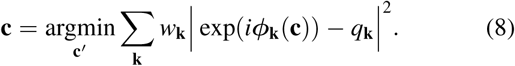

This step requires only seconds of computation time. In addition to making the problem tractable, this separation into two steps also allows us to inspect the phase angles of the per-pixel optima *q*_**k**_ visually and to determine the type of antisymmetrical aberration present in the data set.

After the optimal antisymmetrical aberration coefficients c have been determined, they are used to invert the phase shift of all observed images *X* when a 3D map is being reconstructed from them.

### 2.3. Symmetrical Aberrations

Unlike the antisymmetrical aberrations, the symmetrical ones act on the CTF. In the presence of such aberrations, the CTF no longer consists of strictly elliptical rings of alternating sign, but it can take on a more unusual form. In our experiments, we have specifically observed the ellipses deforming into slightly square-like shapes. In order to estimate the symmetrical aberration, we need to determine the most likely deformations of the CTFs hidden underneath the measured noisy pixels. Since the micrographs in a cryo-EM data set are usually collected under different defoci, it is not sufficient to measure the collective power spectrum of the entire data set – instead, we need to determine one deformation applied to *different* CTFs.

In RELION-3.1, the CTF is defined as:

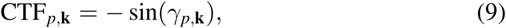

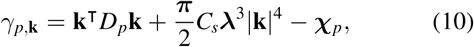

where *D*_*p*_ is the real symmetrical 2 × 2 astigmatic-defocus matrix for particle *p, C*_*s*_ is the spherical aberration of the microscope, λ is the electron wavelength and χ_*p*_ is a constant offset given by the amplitude contrast and the phase shift due to a phase plate (if one is used). We chose this formulation of astigmatism because it is both more concise and also more practical when dealing with anisotropic magnification, as shown in section 2.4. In Appendix A, we define *D*_*p*_ and we show that this is equivalent to the more common formulation (Mindell & Grigorieff, 2003).

We model the deformation of the CTF under symmetrical aberrations by offsetting *γ*:

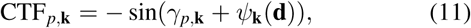

where *ψ*_**k**_(**d**) is modelled using symmetrical Zernike polynomials combined with a set of coefficients **d** ∈ ℝ^*B*^ that describe the aberration:

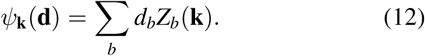

The optimal values of *d*_*b*_ are determined by minimizing another sum of squared differences:

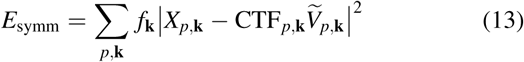

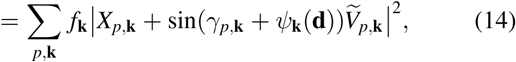

where the predicted complex amplitude 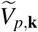 contains the phase shift induced by the antisymmetrical aberration (if it is known):

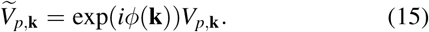

This is again a non-linear equation with a large number of terms. In order to make its minimization tractable, we perform the following substitution:

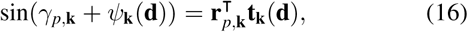

with the known column vector **r**_*p*,**k**_ ∈ ℝ^2^ given by

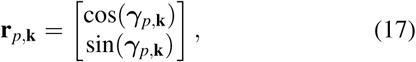

and the unknown **t**_**k**_(**d**) ∈ ℝ^2^ by

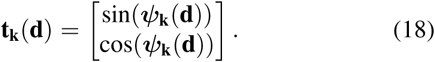

This allows us to transform the one-dimensional non-linear term for each pixel **k** into a two-dimensional linear one:

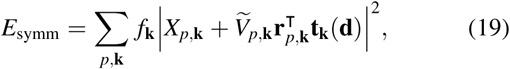

In this form, we can decompose *E*_symm_ into a sum of quadratic functions over all pixels **k**. This is equivalent to the transformation in Eq. 5, only in two real dimensions instead of one complex one:

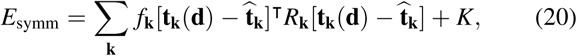

where the real symmetrical 2 × 2 matrix *R*_**k**_ is given by

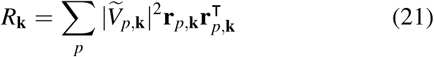

and the corresponding per-pixel optima 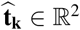 by

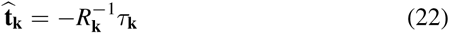

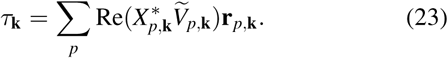

Again, computing *R*_**k**_ and 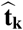 only requires one iteration over the data set, where for each pixel **k**, five numbers need to be updated for each particle *p*: the three distinct elements of *R*_**k**_ (the matrix is symmetrical) and the two of *τ*_**k**_. Once *R*_**k**_ and 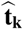 are known, the optimal Zernike coefficients d are determined by minimizing *E*_symm_ in Eq. 20 using the Nelder-Mead downhill simplex algorithm. Analogously to the case of the antisymmetrical aberrations, a visual inspection of the optimal *ψ*_**k**_(**d**) for each pixel allows us to examine the type of aberration without projecting it into the Zernike basis. The CTF phase-shift estimate for pixel **k** is given by 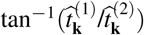, where 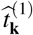 and 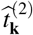 refer to the two components of **t**_**k**_

Once the coefficients d of the symmetrical aberration are known, they are used to correct any CTF that is computed in RELION-3.1.

### 2.4. Anisotropic Magnification

To determine the anisotropy of the magnification, we again compare predicted images to the observed ones. We assume that the 3D reference map *W* has been obtained by averaging views of the particle at in-plane rotation angles drawn from a uniform distribution. This is a realistic assumption, since, unlike the angle between the particle and the ice surface where the particle often shows a preferred orientation, the particle is oblivious to the orientation of the camera pixel-grid. Thus, for a data set of a sufficient size, the anisotropy in the individual images averages out and the resulting reference map depicts an isotropically scaled 3D image of the particle (although the high-frequency information on the periphery of the particle is blurred out by the averaging). We can therefore estimate the anisotropy by determining the optimal deformation that has to be applied to the predicted images in order to best fit the observed ones.

We are only looking for linear distortions of the image. Such a distortion can be equivalently represented in real space or in Fourier space: if the real-space image is distorted by a 2 × 2 matrix *M*, then the corresponding Fourier-space image is distorted by *M*^−1^. We choose to operate in Fourier space since this allows us to determine the deformation of the predicted image without also distorting the CTF. We assume that the CTF is represented correctly in the distorted coordinates because it has been estimated from the original images before the distortion was known.

Formally, we define the complex amplitude *V*_*p*,**k**_(*M*) of the predicted image deformed by a 2 × 2 matrix *M* by:

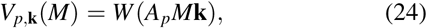

and we aim to determine such a matrix *M* that minimizes:

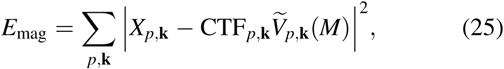

where 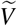 again refers to the phase shifted complex amplitudes as defined in Eq. 15. We are not assuming that *M* is necessarily symmetrical, which allows it to express a skew component in addition to the anisotropic magnification. Such skewing effects are considered by the models commonly used in computer vision applications (Hartley, 1994; Hartley & Zisserman, 2003), but not in cryo-EM. We have decided to model the skew component as well, in case it should manifest in a data set. The expression given in Eq. 25 is yet another sum over a large number of non-linear terms. In order to obtain a sum over squares of linear terms, we first express the deformation by *M* as a set of per-pixel displacements *δ*_**k**_ ∈ ℝ^2^:

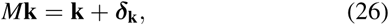

Next, we perform a first-order Taylor expansion of *W* around *A*_*p*_**k**. We know that this linear approximation of *W* is reasonable for all frequencies **k** at which the reference map contains any information, because the displacements *δ*_**k**_ are smaller than one voxel there. If they were significantly larger, then they would prevent a successful computation of the complex amplitudes of the reference map at those frequencies. The linear approximation is given as follows:

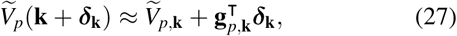

where the gradient **g**_*p*,**k**_ ∈ ℂ^2^ is a column vector that is computed by forward projecting the 3D gradient of *W* (which is given by the linear interpolation):

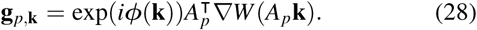

It is essential to compute **g**_*p*,**k**_ in this way, since computing it numerically from the already projected image *V*_*p*,**k**_ would lead to a systematic underestimation of the gradient (due to the interpolation) and thus to a systematic overestimation of the displacement. Note also that the change in *ϕ*(**k**) as a result of the displacement is being neglected. This is due to the fact that the phase shift, like the CTF, has also been computed from the distorted images, so that we can assume it to be given correctly in the distorted coordinates.

Using the terms transformed in this way, the sum of squared errors can be approximated by:

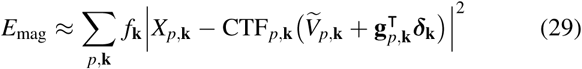

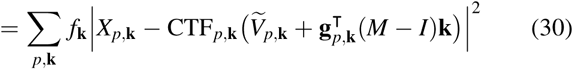

This corresponds to two linear systems of equations to be solved in a least-squares sense, either for the per-pixel displacements *δ*_**k**_ (Eq. 29) or for the global deformation matrix *M* (Eq. 30). Analogously to the aberrations methods, we solve for both. Knowing the per-pixel solutions again allows us to confirm visually whether the observed deformations are consistent with a linear distortion: if they are, then the per-pixel displacements *δ*_**k**_ will follow a linear function of **k**.

The optimal displacements 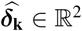 are equal to:

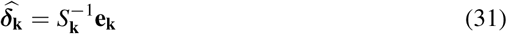

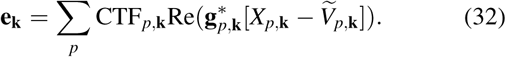

with the real symmetrical 2 × 2 matrix *S*_**k**_ given by:

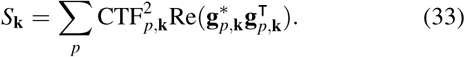

Note that this is equivalent to treating the real and imaginary components of Eq. 29 as separate equations, since Re(z\**w*) = Re(*z*)Re(*w*) + Im(*z*)Im(*w*) for all *z, w* ∈ ℂ. Analogously to the estimation of the symmetrical aberrations, *S*_**k**_ and **e**_**k**_ are computed in one iteration by accumulating five numbers for each pixel **k** over the entire data set.

The optimal 2 × 2 deformation matrix *M* is determined by first reshaping it into a column vector **m** ∈ ℝ^4^:

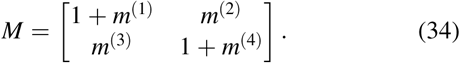

The expression in Eq. 30 can then be written as

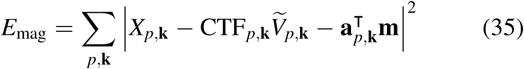

with the column vector **a**_*p*,**k**_ ∈ ℂ^4^ given by

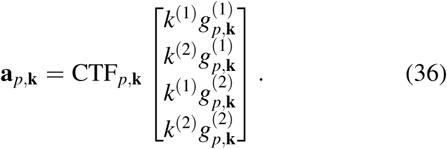

We can now compute the optimal **m**:

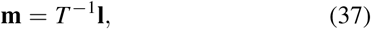

where the real symmetrical 4 × 4 matrix *T* and the column vector **l** ∈ ℝ^4^ are equal to

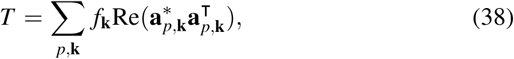

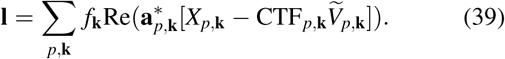

There is no need to compute *T* and **l** explicitly by iterating over all particles *p* again, since all the necessary sums are already available as part of *S*_**k**_ and **e**_**k**_. Instead, we only need to sum up the corresponding values over all pixels **k**. This is shown in Appendix B.

In order to correct for the anisotropy after *M* has been estimated, we never resample the observed images. When we compute a 3D map from a set of observed images, we do so by inserting 2D slices into the 3D Fourier-space volume. Since this process requires the insertion of 2D pixels at fractional 3D coordinates (and thus interpolation), we can avoid any additional resampling of the observed images by instead inserting pixel **k** into the 3D map at position *A*_*p*_*M***k** instead of at *A*_*p*_**k**. Analogously, if the methods described in 2.2 and 2.3 are applied after the distortion matrix *M* is known, then the predicted images are generated by reading the complex amplitude from *W* at 3D position *A*_*p*_*M***k**. This has been omitted from the description of those methods to aid readability.

Furthermore, when dealing with anisotropic magnification in RELION, we have chosen to always define the CTF in the undistorted 2D coordinates. The primary motivation behind this is the assumption that the spherical aberration (second summand in Eq. 10) should only be radially symmetrical if the image is not distorted. For this reason, once the distortion matrix *M* is known, we need to transform the astigmatic-defocus matrix *D* into the new undistorted coordinate system. This is done by conjugating *D* under *M*^−1^:

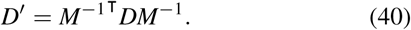

When a CTF value is computed after this transformation has been performed, it is always computed as CTF(*M***k**) instead of as CTF(**k**).

The Zernike polynomials that are used as a basis for the symmetrical and antisymmetrical aberrations are also defined in the undistorted coordinates, i.e. the Zernike polynomials are also evaluated at *Z*_*b*_(*M***k**). Note that these coefficients cannot be trivially corrected after estimating *M*. Instead, we propose that the aberrations be estimated only after *M* is known. In severe cases, a better estimate of *M* can be obtained by repeating the magnification refinement after determining optimal defocus and astigmatism estimates using the initial estimate of *M*. We illustrate this scenario on a synthetic example in section 3.4.

### 2.5. Implementation Details

The three methods described above need to be applied to a large number of particles in order to obtain a reliable estimate. Nevertheless, we allow the three effects to vary within a data set in RELION-3.1. To facilitate this, we have introduced the concept of *optics groups*: partitions of the particle set that share the same optical properties, such as the voltage or pixel size (or the aberrations and the magnification matrix). As of RELION-3.1, those optical properties are allowed to vary between optics groups, while particles from different groups can still be refined together. This makes it possible to merge data sets collected on different microscopes with different magnifications and aberrations without the need to resample the images. The anisotropic magnification refinement can then be used to measure the relative magnification between the optics groups, by refining their magnification against a common reference map.

Since most of the optical properties of a particle are now defined through the optics group to which it belongs, each particle STAR file written out by RELION-3.1 now contains *two* tables: one listing the optics groups and one listing the particles. The particles table is equivalent to the old one, except that certain optical properties are no longer listed. Those are typically the voltage, the pixel and image sizes, the spherical aberration and the amplitude contrast, and they are instead specified in the optics groups list. This reduces the overall file size, and it makes manual editing of those properties easier.

A number of other optical properties are still stored in the particles list, allowing for different values for different particles in the same group. These properties make up the per-particle part of the symmetrical aberration, i.e. the coefficient *γ*_*p*,**k**_ in Eq. 10. The specific parameters that can vary per particle are the following: the phase shift, defocus, astigmatism, the spherical aberration and the B-factor envelope.

We have developed a new CTF refinement program that considers all particles in a given micrograph, and locally optimises all of the above five parameters, while each parameter can be modelled either per particle, per micrograph or remain fixed. The program then uses the L-BFGS algorithm (Liu & Nocedal, 1989) to find the least-squares optimal parameter configuration given all the particles in the micrograph. This allows the user to find for example the most likely phase shift of a micrograph while simultaneously finding the most likely defocus value of each particle in it.

Note that the terms *defocus* and *astigmatism* above refer specifically to *δz* (defocus) and *a*_1_ and *a*_2_ (astigmatism), where the astigmatic defocus matrix *D*_*p*_ of particle *p* in Eq. 10 is composed as follows:

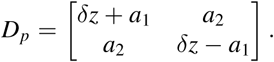

As an example, this would allow the defocus to be expressed per particle, by allocating a separate *δz* for each particle, while the astigmatism could be estimated per micrograph by requiring *a*_1_ and *a*_2_ to be identical for all particles.

Like the astigmatism, the B-factor envelope is also a two dimensional parameter, and it consists of a scale factor *S* and the *B* factor itself. It corresponds to a Gaussian envelope over the CTF (given by 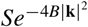) and it provides a means of weighting different particles against each other. Specifically, a greater *B* factor means that the particle will contribute less to the higher frequencies of the reconstruction. Although *B* factors on the CTF have been available in earlier releases of RELION, the method to estimate them is new in version 3.1.

## 3. Results

To validate our methods and to illustrate their usefulness, we describe four experiments using publicly available data sets. First, we assess aberration correction on two data sets that were collected on a 200 keV Thermo Fischer Talos Arctica microscope. Second, we illustrate a limitation of our method for modelling aberrations using a data set that was collected on a 300 keV Thermo Fischer Titan Krios microscope with a Volta phase-plate with defocus (Danev *et al.*, 2017). Third, we apply our methods to one of the highest-resolution cryo-EM structures published so far, collected on a Titan Krios without a phase plate. Finally, we determine the precision to which the magnification matrix *M* can be recovered in a controlled experiment, using artificially distorted images, again from a Titan Krios microscope.

### 3.1. Aberrations Experiment at 200 keV

We re-processed two publicly available data sets: one on the rabbit muscle aldolase (EMPIAR-10181), the other on the *T. acidophilum* 20S proteasome (EMPIAR-10185). Both data sets were collected on the same 200 keV Talos Arctica microscope, which was equipped with a Gatan K2 Summit direct electron camera. At the time of the original publication (Herzik Jr *et al.*, 2017), the aldolase could be reconstructed to 2.6 Å the proteasome to 3.1 Å using RELION-2.0.

We picked 159,352 particles for the aldolase data set, and 74,722 for the proteasome. For both data sets, we performed five steps and measured the resolution at each step. First, we refined the particles without considering the aberrations. The resulting 3D maps were then used to perform an initial CTF refinement in which the per-particle defoci and the aberrations were estimated. The particles were then subjected to Bayesian polishing (Zivanov *et al.*, 2019), followed by another iteration of CTF refinement. In order to disentangle the effects of improved Bayesian polishing from the aberration correction, we also performed a refinement with the same polished particles, but assuming all aberrations to be zero. We measured the Fourier-shell correlation (FSC) between the two independent half sets and against known reference structures (PDB-1ZAH and PDB-6BDF, respectively (St-Jean *et al.*, 2005; Campbell *et al.*, 2015)). The plots are shown in Fig. 1, and the resolutions measured by the half-set method, using a threshold of 0.143, in Table 1. Plots of the aberration estimates are shown in Fig. 2.

**Table 1.** Half-set resolutions obtained at different stages of our processing pipeline in the aberration experiment on aldolase and 20S proteasome at 200 keV.

|  | aldolase | proteasome |
| --- | --- | --- |
| initial | 2.7 Å | 3.2 Å |
| first CTF refinement | 2.4 Å | 2.5 Å |
| Bayesian polishing | 2.3 Å | 2.3 Å |
| **second CTF refinement** | **2.1 Å** | **2.3 Å** |
| no aberrations | 2.5 Å | 3.1 Å |

**Figure 1.**
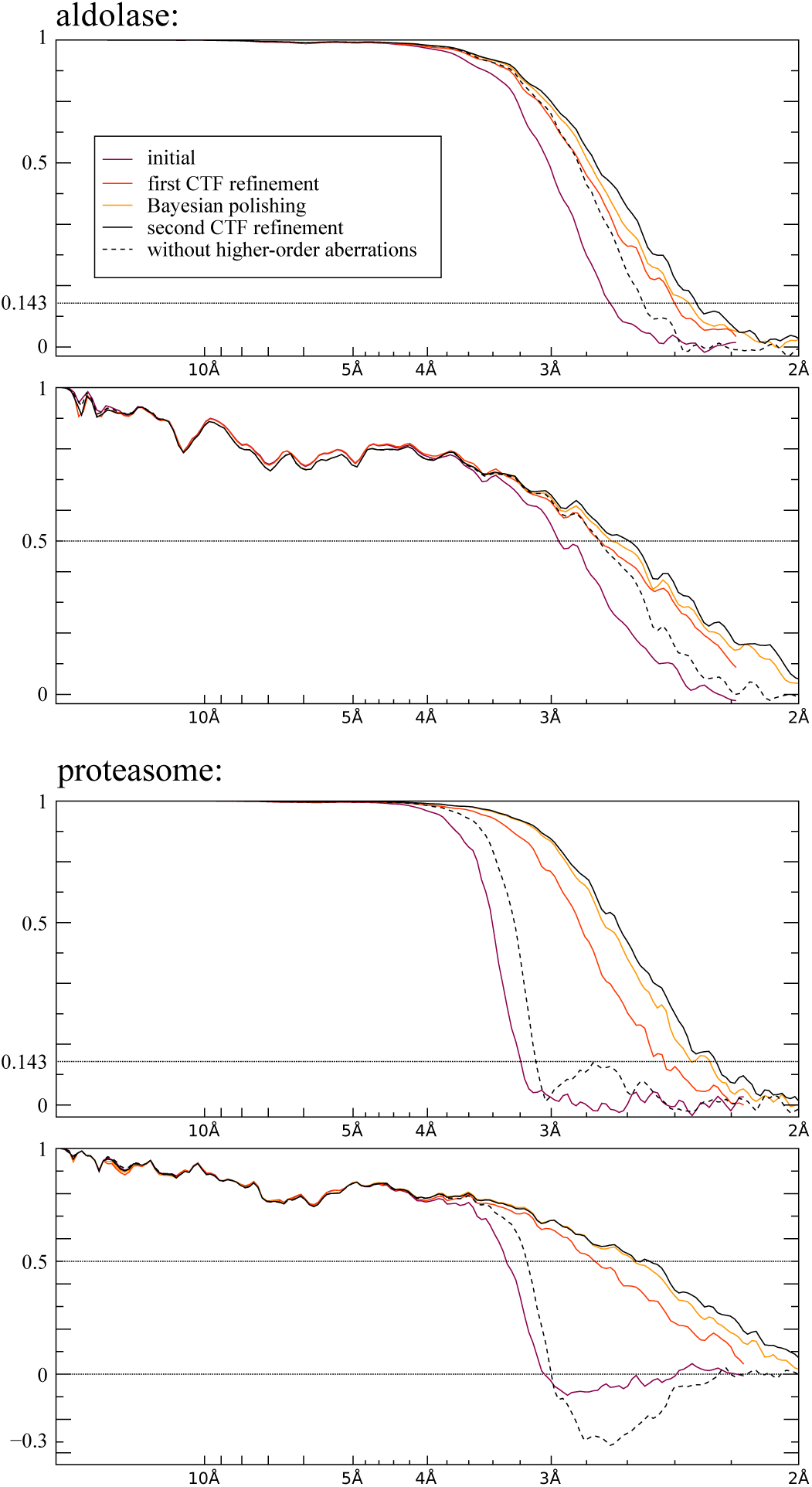
FSC plots from the aberration experiments on aldolase and 20S proteasome at 200 keV. The top plot shows the half-set FSC and the bottom one the FSC against the respective reference structure (see text for details). Note that estimating the aberrations during the initial CTF refinement already produces a significant increase in resolution (red line). It also allows for more effective Bayesian polishing and defocus refinement, increasing the resolution further (black line). Neglecting the aberrations while keeping the remainder of the parameters the same (dashed line) allows us to isolate the effects of aberration correction. For the proteasome, it also exposes a slight (false) positive peak in the half-set FSC around 2.7Å which corresponds to a negative peak in the reference FSC. This indicates that the phases of the complex amplitudes of the 3D map are, on average, flipped at that frequency band due to the strong aberrations.

**Figure 2.**
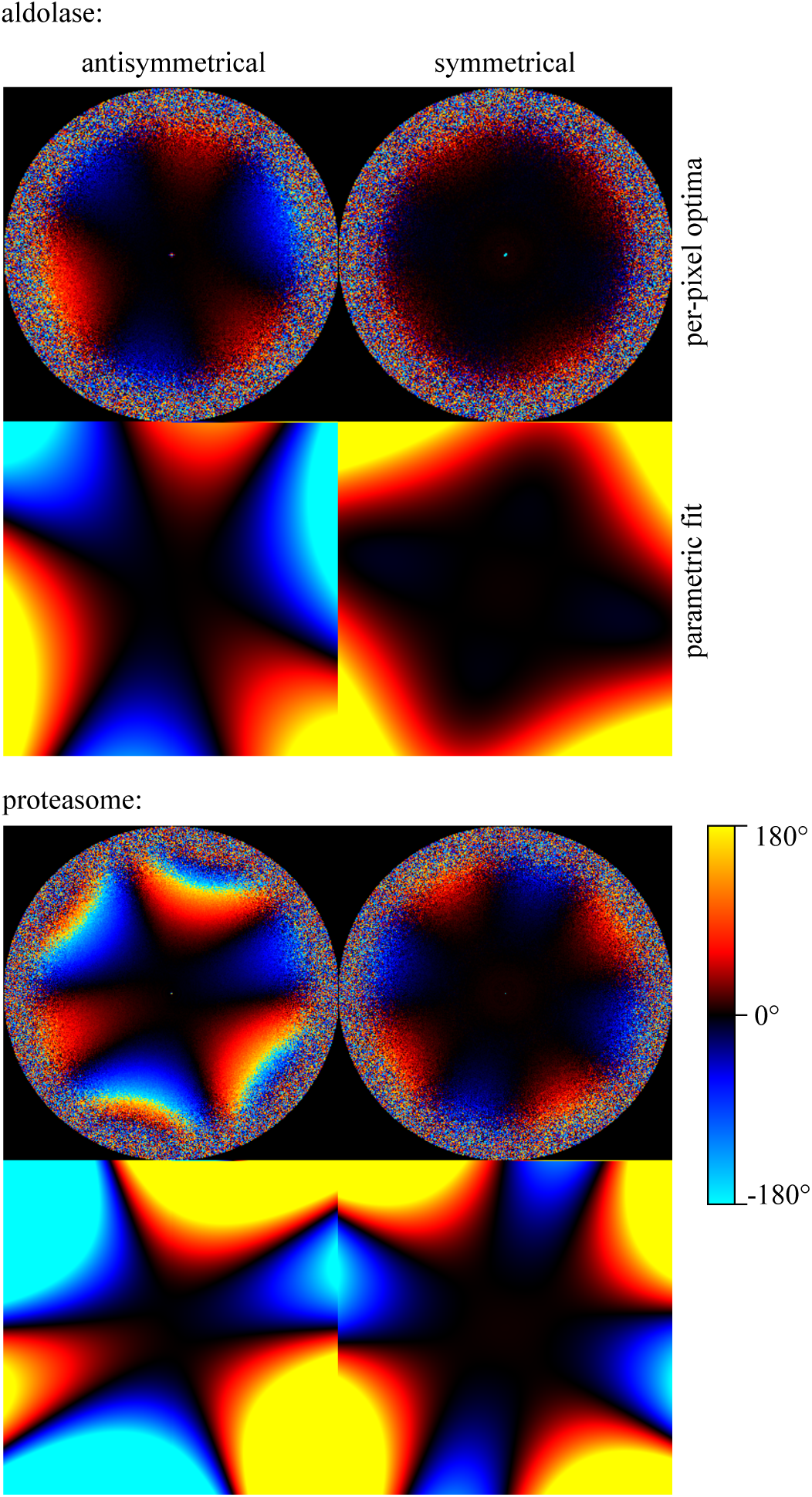
Antisymmetrical and symmetrical aberration experiments on aldolase and 20S proteasome at 200 keV. The upper image of each pair shows the independent phase-angle estimates for each pixel, while the lower shows the parametric fit using Zernike polynomials. These types of aberrations are referred to as *trefoil* or *three-fold astigmatism* (left) and *four-fold astigmatism* (right). The proteasome trefoil exceeds 180° at the very high frequencies, so the sign in the per-pixel plot wraps around. This has no impact on the parametric fit.

Fig. 2 indicates that both data sets exhibit antisymmetrical as well as symmetrical aberrations. For both data sets, the shapes of both types of aberrations are well visible in the per-pixel plots, and the parametric Zernike fits capture those shapes well. The antisymmetrical aberrations correspond to a trefoil (or three-fold astigmatism) combined with a slight axial coma and they are more pronounced than the symmetrical ones. The trefoil is visible as three alternating areas of positive and negative phase difference, with approximate three-fold symmetry, in the images for the antisymmetrical aberration estimation (on the left in Fig. 2). The axial coma breaks the three-fold symmetry, by making one side of the image more positive and the opposite side more negative. The apparent four-fold symmetry in the images for the symmetrical aberrations (on the right in Fig. 2) correspond to four-fold astigmatism and are strongest for the proteasome data set. The proteasome also shows the stronger antisymmetrical aberrations, which even exceed 180° at the higher frequencies. Note that because the per-pixel plots show the *phase angle* of 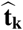 from Eq. 20, they wrap around once they reach 180°. This has no effect on the estimation of the parameters, however, since 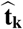 itself, which is a 2D point on a circle, is used in the optimisation and not its phase angle.

The FSC plots (Fig. 1) indicate that aberration correction leads to higher resolution, as measured by both the FSC between independently refined half-maps and the FSC against the reference structure. Comparing the result of the second CTF refinement and its equivalent run without aberration correction (lower two lines in Table 1), the resolution increased from 2.5 Å to 2.1 Å for the aldolase data set, and from 3.1 Å to 2.3 Å for the proteasome. In addition, aberration correction also allows for more effective Bayesian polishing and defocus estimation, which is the reason for performing the CTF refinement twice.

### 3.2. Phase-Plate Experiment

We also analysed a second data set on a *T. acidophilum* 20S proteasome (EMPIAR-10078). This data set was collected using a Volta phase-plate (VPP) (Danev *et al.*, 2017) under defocus. We picked 138,080 particles and processed them analogously to the previous experiment, except that the CTF refinement now included the estimation of anisotropic magnification. The estimated aberrations are shown in Fig. 4 and the FSCs in Fig. 6.

**Figure 3.**
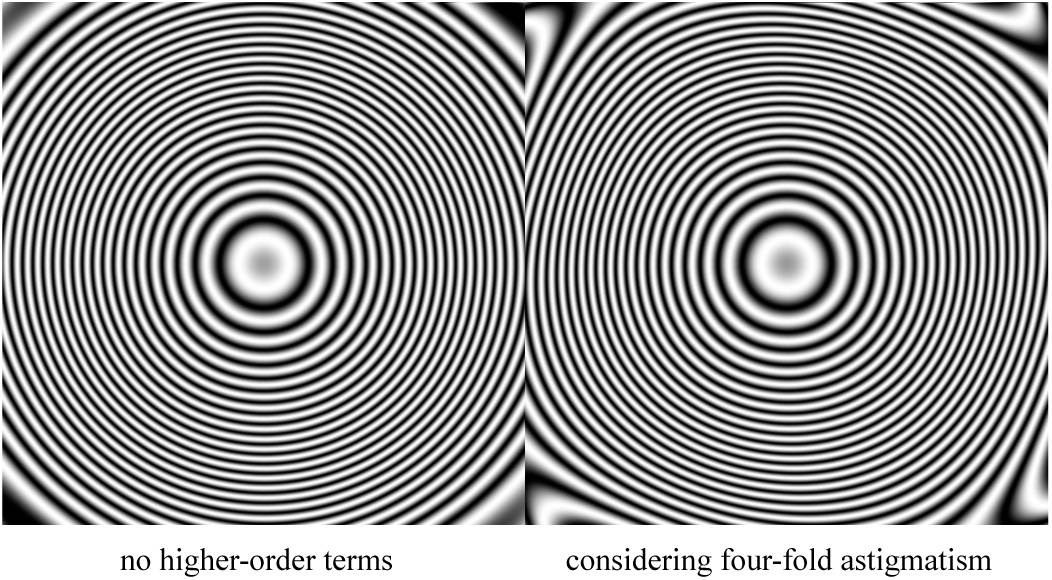
Effects of the symmetrical aberrations on the CTF of the 20S proteasome as part of the aberration experiment at 200 keV. The image on the left shows a CTF expressed by the traditional model, while the one on the right shows the fit of our new model which considers higher-order symmetrical aberrations. Note that the slightly square-like shape that arises from four-fold astigmatism cannot be expressed by the traditional model. The aberrations correspond to the bottom right image in Fig. 2

**Figure 4.**
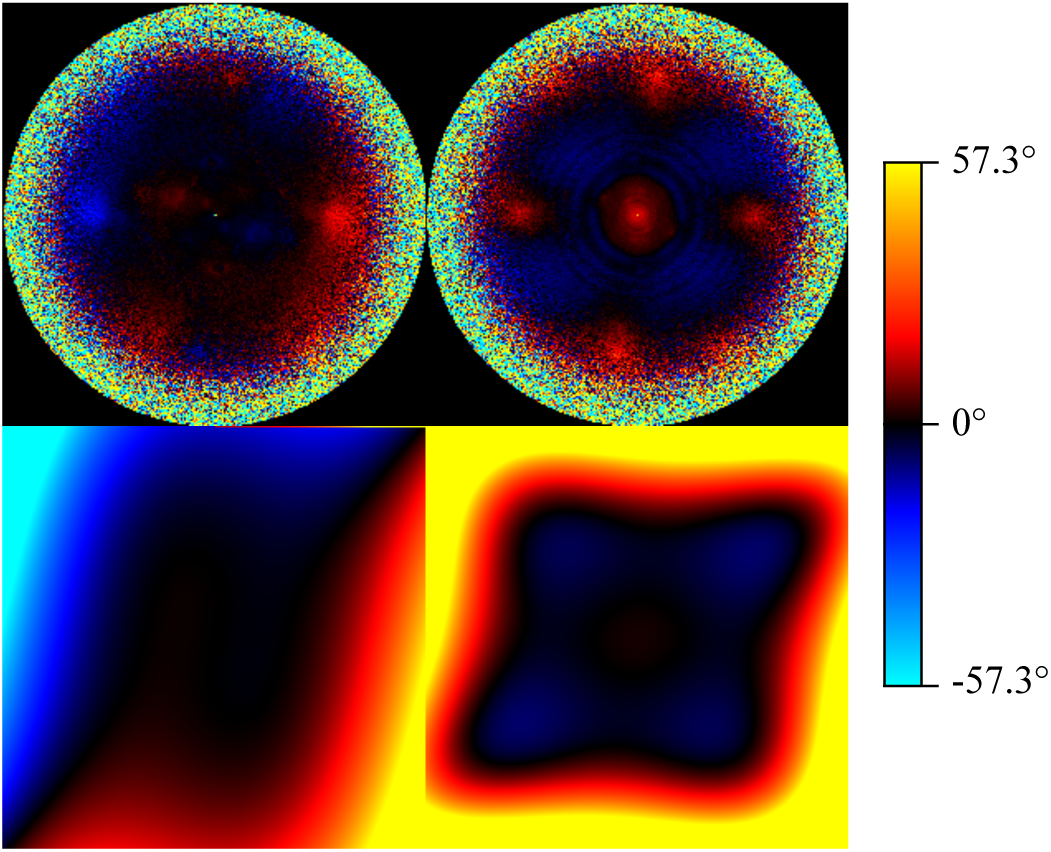
Antisymmetrical (left) and symmetrical (right) aberrations measured on the phase plate data set. The upper image shows independent per-pixel estimates and the lower the parametric fits. Note the four afterimages of previously used phase-plate spots in the upper right image. They cannot be represented by our model.

**Figure 5.**
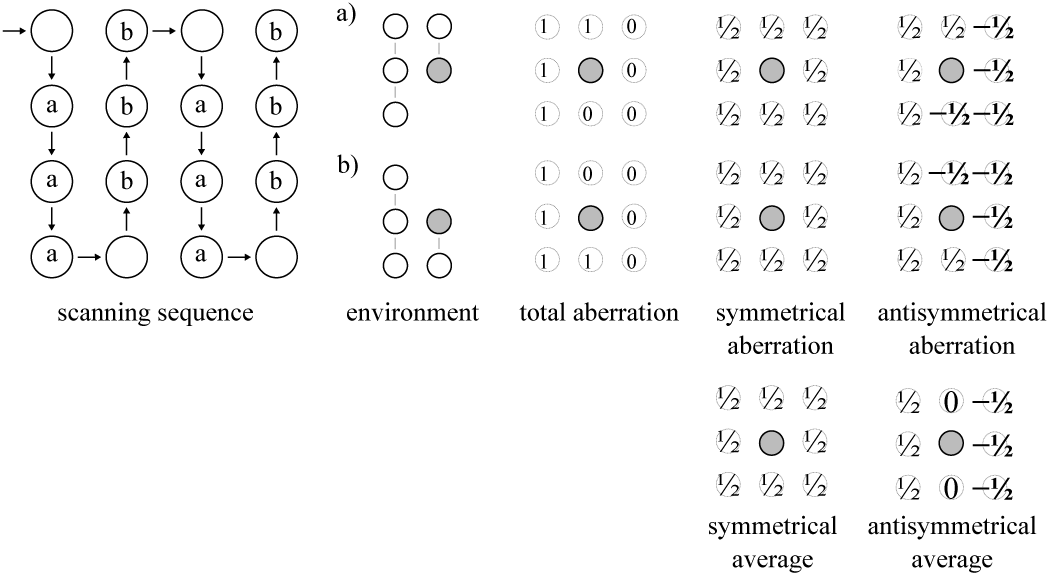
Our interpretation of the aberration plots in Fig. 4. The presence of all four neighbouring spots in the symmetrical plot, together with the absence of the vertical neighbours from the antisymmetrical plot, indicates that the VPP spots were scanned in a vertically alternating and horizontally unidirectional sense, as shown in the first image. This partitions a majority of the spots into two classes, *a* and *b*, in which the direct vertical neighbour is located on opposite sides. The total phase shift induced by the neighboring spots is decomposed into an antisymmetrical and a symmetrical part. Both of them are averaged over particles from both classes during estimation, so the vertical neighbor partially cancels out in the antisymmetrical plot, but not in the symmetrical one.

**Figure 6.**
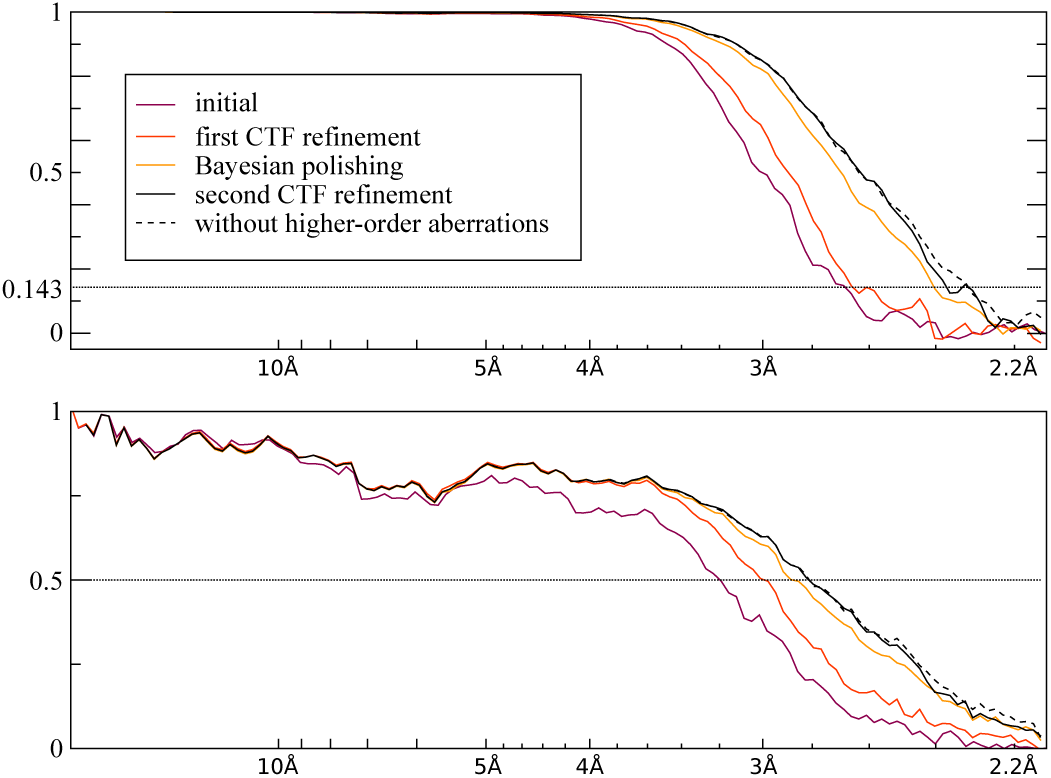
Half-set (top) and reference (bottom) FSC plots for the phase plate data set. The reference structure used was again PDB-6BDF. Note that considering the aberrations does not improve the resolution, since these types of aberrations cannot be expressed by our model. Nevertheless, the CTF refinement does improve the resolution due to the new micrograph-global defocus and phase-shift estimation and due to considering the slightly anisotropic magnification.

The purpose of a VPP is to shift the phase of the unscattered beam in order to increase the contrast against the scattered beam. This is accomplished by placing a heated film of amorphous carbon (the VPP) at the back-focal plane of the microscope and letting the electron beam pass through it after it has been scattered by the specimen. The central, unscattered beam – which exhibits much greater intensity than the unscattered components – then spontaneously creates a spot of negative electric potential on the VPP (Danev *et al.*, 2014). It is this spot which then causes the phase shift in the unscattered beam. After being used for a certain amount of time, the spot charges up even more and develops imperfections. At that point, the user will typically switch to a different position on the carbon film. The charge at the previous position will decay, although some charge may remain for an extended period. If the VPP is shifted by an insufficient distance, the old spot will reside in a position traversed by scattered rays corresponding to some higher image frequency. We hypothesize that we can observe these spots in our symmetrical aberration plots.

The symmetrical plots show a positive phase shift at the center of frequency space (Fig. 4). We hypothesize that this spot is caused by the size of the charge built-up at the currently used position on the phase-plate (Danev & Baumeister, 2016). Moreover, this plot shows four additional spots at higher spatial frequencies. We hypothesize that these may arise from residual charges on previously used phase-plate positions. These charges would then interfere with the diffracted rays at higher spatial frequency from the current position, resulting in the observed spots in the aberration image. The absence of the vertical neighbor-spots from the antisymmetrical plot suggests that the spots were scanned in a vertically alternating but horizontally unidirectional sense. This is illustrated in Fig. 5.

Because these types of aberrations do not satisfy our smoothness assumptions, they cannot be modelled well using a small number of Zernike basis polynomials. Although increasing the number of Zernike polynomials would in principle allow expressing any arbitrary aberration function, it also decreases the system’s ability to extrapolate the aberration into the unseen high-frequency regions. As a consequence, our aberration model cannot be used to neutralise the effects of the phaseplate positions, which is confirmed by the FSC plots in Fig. 6. In practice, this problem can be avoided experimentally, by spacing the phase plate positions further apart and thus arbitrarily increasing the affected frequencies.

The estimated magnification anisotropy for this data set is relatively weak. The final magnification matrix *M* we recovered was:

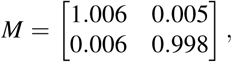

which corresponds to 1.35% anisotropy along two perpendicular axes rotated by 66°.

### 3.3. High-Resolution Experiment

We applied our methods to a mouse heavy-chain apoferritin data set (EMPIAR-10216) collected on a 300 keV Titan Krios fitted with a Falcon 3 camera. At the time of its publication, the particle could be reconstructed to a resolution of 1.62 Å using RELION-3.0 (Danev *et al.*, 2019). This data set thus offers us a means to examine the effects of higher-order aberrations and anisotropic magnification at higher resolutions.

We compared the following three reconstructions. First, the original, publicly available map. Since it had been estimated using RELION-3.0, only the effects of beam tilt could be corrected for, but none of the other high-order aberrations or anisotropic magnification. Second, the aberrations alone: for this, we proceeded from the previous refinement, and we first estimated the higher order aberrations and then, simultaneously, per-particle defoci and per-micrograph astigmatism. Third, we performed the same procedure, but only after first estimating the anisotropic magnification. For the third case, the entire procedure was repeated after a round of refinement. For all three cases, we calculated the FSC between the independently refined half-maps and the FSC against an atomic model, PDB-6S61, that was built in an independently reconstructed cryo-EM map of mouse apoferritin at a resolution of 1.84 Å. In the absence of a higher-resolution reference structure, comparison with PDB-6S61 relies on the assumption that the geometrical restraints applied during atomic modelling resulted in predictive power at resolutions beyond 1.84 Å. We used the same mask as in the original publication for correction of the solvent-flattening effects on the FSC between the independent half-maps, and we used the same set of 147,637 particles throughout.

The aberration plots in Fig. 7 show that this data set exhibits a trefoil aberration and faint four-fold astigmatism. In the magnification plot in Fig. 8, we can see a clear linear relationship between the displacement of each pixel **k** and its coordinates. This indicates that the measured displacements stem from a linearly distorted image and that the implied distortion is a horizontal dilation and a vertical compression. This is consistent with anisotropic magnification, since the average magnification has to be close to 1 because the reference map itself has been obtained from the same images under random in-plane angles. The smoothness of the per-pixel plot suggest that the large number of particles allows us to measure the small amount of anisotropy reliably. The magnification matrix we estimated was:

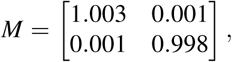

which corresponds to 0.54% anisotropy. As can be seen in the FSC curves in Fig. 9, considering either of these effects is beneficial, while considering both yields a resolution of 1.57 Å, an improvement of three shells over the reconstruction obtained using RELION-3.0.

**Figure 7.**
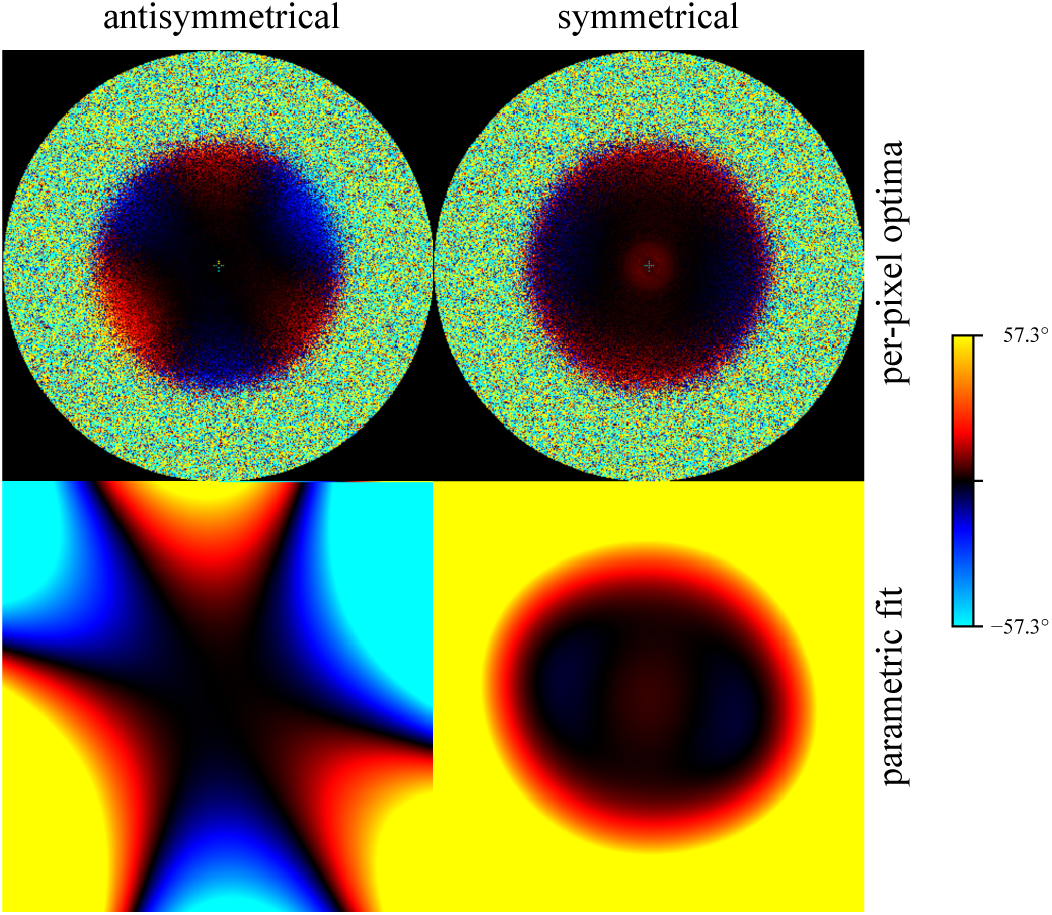
Higher-order aberrations measured on the high-resolution mouse apoferritin data set. The antisymmetrical plot (left) shows a significant trefoil aberration, while the symmetrical plot (right) shows a faint four-fold astigmatism. Although the aberrations are comparatively weak, they are clearly measurable and considering them does lead to a small improvement in resolution (see Fig. 9).

**Figure 8.**
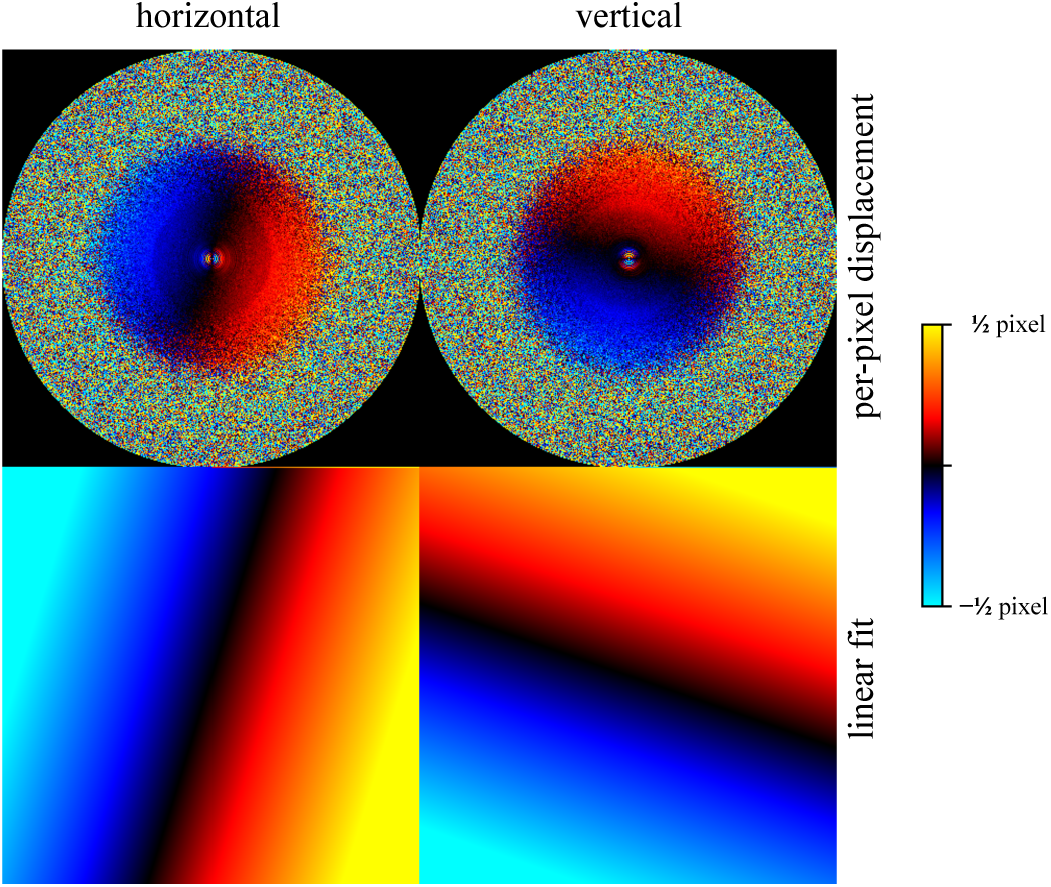
Anisotropic magnification plots for the high-resolution mouse apoferritin data set. The top row shows the estimated displacement for each pixel (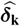 in Eq. 31) while the bottom row shows the displacement corresponding to the estimated magnification matrix *M* (i.e., *M***k** − **k**). Note that the per-pixel estimates follow a linear relationship, indicating that the displacements are indeed caused by a linear transformation of the image. The horizontal coordinate is defined as increasing to the right and the vertical as increasing downward, so the two plots indicate a horizontal dilation and a vertical compression.

**Figure 9.**
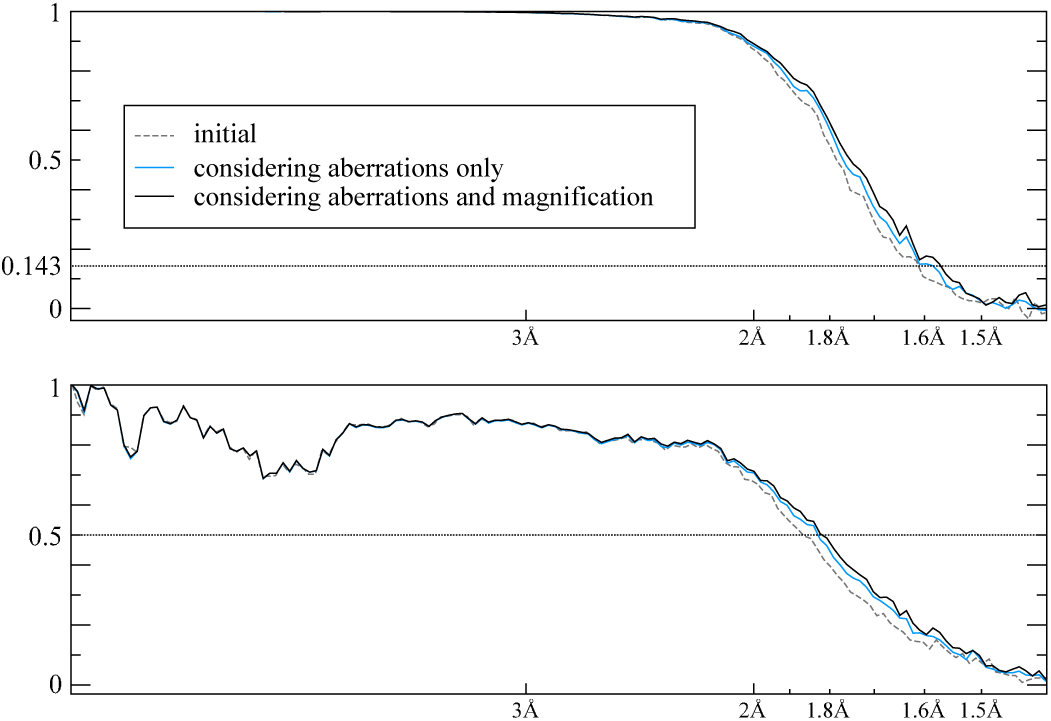
Half-set (top) and reference (bottom) FSC plots for the high-resolution mouse apoferritin data set. Considering the anisotropic magnification (black line) produces a further improvement in terms of resolution beyond what is attainable by considering the aberrations alone (blue line). The reference map used was PDB-6S61, another publicly availably cryo-EM structure. The resolution indicated by the bottom plot is limited by the fact that the resolution of the reference structure is only 1.84Å.

### 3.4. Simulated Anisotropic Magnification Experiment

To measure the performance of our anisotropic magnification estimation procedure in the presence of a larger amount of anisotropy, we also performed an experiment on synthetic data. For this experiment, we used a small subset (9, 487 particles from 29 movies) taken from a human apoferritin data set (EMPIAR-10200), which we had processed before (Zivanov *et al.*, 2018). We distorted the micrographs by applying a known anisotropic magnification using MotionCor2 (Zheng *et al.*, 2017). The relative scales applied to the images were 0.95 and 1.05, respectively, along two perpendicular axes rotated at a 20° angle. In this process, about 4% of the particles were mapped outside the images, so the number of distorted particles is slightly smaller, 9,093.

We then performed 4 rounds of refinement on particle images extracted from the distorted micrographs in order to recover the anisotropic magnification. Each round consisted of a CTF refinement followed by an autorefinement. The CTF refinement itself was performed twice each time, once to estimate the anisotropy, and then again to determine the per-particle defoci and per-micrograph astigmatism. The FSC curves for the different rounds can be seen in Fig. 10. We observe that the FSC approaches that of the undistorted particles already after the second round. In the first round, the initial 3D reference map is not precise enough to allow for a reliable recovery of anisotropy.

**Figure 10.**
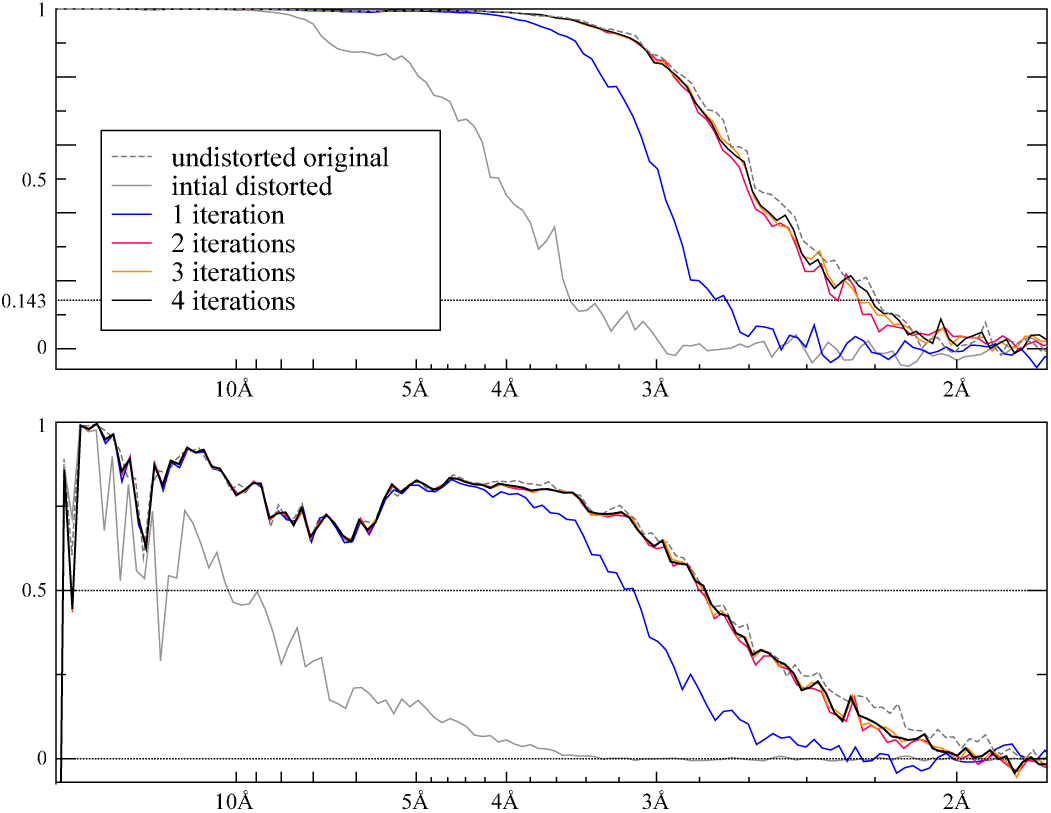
Half-set (top) and reference (bottom) FSC plots for the simulated anisotropic magnification experiment on human apoferritin. The reference structure used was PDB-5N27 (Ferraro *et al.*, 2017). From the second iteration onward, the curves lie close to their final position. Note that the resolution of the undistorted reconstruction cannot be reached by the distorted ones, since particles have been lost along the way and since the image pixels have been degraded by resampling.

The magnification matrix M recovered in the final round looks as follows:

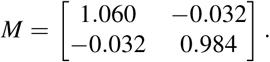

It corresponds to the relative scales of 0.951 and 1.049, respectively, along two perpendicular axes rotated by 19.939°, although it also contains an additional uniform scaling by a factor of 1.022. The uniform scaling factor has no influence on the refinement, but it does change the pixel size of the resulting map. We therefore note that caution must be taken to either enforce the product of the two relative scales to be 1, or to otherwise calibrate the pixel size of the map against an external reference.

This experiment shows that the anisotropy of the magnification can be estimated to 3 significant digits, even from a relatively small number of particles. Since the estimate arises from adding up contributions from all particles, the precision increases with their number.

## 4. Discussion

Although we previously described a method to estimate and correct for beam-tilt-induced axial coma (Zivanov *et al.*, 2019), no methods to detect and correct for higher-order optical aberrations were available until now. It is therefore not yet clear how often these aberrations are a limiting factor in cryo-EM structure determination of biological macromolecules. The observation that we have already encountered several examples of strong three-and four-fold astigmatism on two different types of microscopes suggests that these aberrations may be relatively common.

Our results with the aldolase and 20S proteasome data sets illustrate than when antisymmetrical and/or symmetrical aberrations are present in the data, our methods lead to an important increase in achievable resolution. Both aldolase and the 20S proteasome could be considered as “easy” targets from cryo-EM structure determination – they have both been used to test the performance of cryo-EM hardware and software, e.g. (Li *et al.*, 2013; Danev & Baumeister, 2016; Herzik Jr *et al.*, 2017; Kim *et al.*, 2018). However, our methods are not limited to standard test samples, and have already been used to obtain biological insights on much more challenging data. Images on brain-derived tau filaments from an ex-professional American football player with chronic traumatic encephalopathy that we recorded on a 300keV Titan Krios microscope showed severe three-and four-fold astigmatism. Correction for these aberrations led to an increase in resolution from 2.7 Å to 2.3 Å, which allowed visualisation of alternative side chain conformations and of ordered water molecules inside the amyloid filaments (Falcon *et al.*, 2019).

Titan Krios microscopes come equipped with hexapole lenses that can be tuned to correct for three-fold astigmatism, although this operation is typically only performed by engineers. The Titan Krios microscope that was used to image the tau filaments from the American football player is part of the UK national cryo-EM facility at Diamond (Clare *et al.*, 2017). After measuring the severity of the aberrations, its lenses were readjusted, and no higher-order aberrations have been detected on it since (Peijun Zhang, personal communication). Talos Arctica microscopes do not have lenses to correct for trefoil, and the microscope that was used to collect the aldolase and the 20S proteasome data sets at the Scripps Research Institute continues to yield data sets with fluctuating amounts of aberrations (Gabriel Lander, personal communication). Until the source of these aberrations are determined or better understood, the corrections proposed here will be important for processing of data acquired on these microscopes.

To what extent higher-order aberrations are limiting will depend on the amount of three-and four-fold astigmatism, as well as on the target resolution of the reconstruction. We have only observed noticeable increases in resolution for data sets that yielded reconstructions with resolutions beyond 3.0-3.5 Å before the aberration correction. However, the effects of the aberrations are more pronounced for lower-energy electrons. Therefore, our methods may become particularly relevant for data from 100 keV microscopes, the development of which is envisioned to yield better images for thin specimens and to bring down the elevated costs of modern cryo-EM structure determination (Peet *et al.*, 2019; Naydenova *et al.*, 2019).

The effects of anisotropic magnification on cryo-EM structure determination of biological samples have been described previously (Grant & Grigorieff, 2015; Zhao *et al.*, 2015). This has resulted in awareness of this problem in the field, and several methods to estimate and correct for the presence of anisotropic magnification. However, measuring anisotropic magnification from the difference between defoci in two perpendicular directions depend on the astigmatism being small compared to the anisotropy in the magnification, while measuring the ellipticity of rings in power spectra from multicrystalline test specimens requires additional experiments. By accumulating the differences between reference projections with high signal-to-noise ratios and the particle images of an entire data set, our method has the potential to detect smaller deviations than the existing methods. Such small deviations may be imperceptible at lower spatial frequencies, but will become increasingly important at higher spatial frequencies, as we demonstrate for the mouse apoferritin data set. In addition, our method is, in principle, capable of detecting and modeling skew components in the magnification.

In addition to modeling anisotropic magnification, our method can also be used for the combination of different data sets with unknown relative magnifications. In cryo-EM imaging, the magnification is often not exactly known. Again, it is possible to accurately measure the magnification using crystalline test specimens with known diffraction geometry, but in practice, errors of up to a few percent in the nominal pixel size are often observed. When processing data from a single data set, such errors can be absorbed, to some extent, in the defoci values. Therefore, a small error in pixel size only becomes a problem at the atomic modeling stage, where it leads to overall contracted or expanded models with bad stereochemistry. (Please note that this is no longer true at high spatial frequencies due to the absolute value of the *C*_*s*_; e.g. beyond 2.5 Å for non-*C*_*s*_-corrected 300kV microscopes.) When data sets from different sessions are combined, however, errors in their relative magnification will affect the 3D reconstruction at much lower resolutions. Our method can directly be used to correct for such errors. In addition, to provide further convenience, our new implementation allows for the combination of particle images with different pixel and box sizes into a single refinement. The performance of our methods under these conditions remains to be illustrated.

Our results illustrate that antisymmetrical and symmetrical aberrations, as well as anisotropic magnification, can be accurately estimated and modelled *a posteriori* from a set of noisy projection images of biological macromolecules. No additional test samples or experiments at the microscope are necessary; all that is needed is a 3D reconstruction of sufficient resolution that the optical effects become noticeable. Our methods could therefore in principle be used in a “shoot first, ask questions later” type of approach, where speed of image acquisition is prioritised over exhaustively optimising the microscope’s settings. In this context, we caution that while the boundaries of applicability of our methods remain to be explored, it may be better to reserve their use for unexpected effects in data from otherwise carefully conducted experiments.

We thank Rado Danev for providing polished particles for the data set in EMPIAR-10216, and Jake Grimmett and Toby Darling for assistance with high-performance computing. This work was funded by the UK Medical Research Council (MC UP A025 1013 to SHWS), the Japan Society for the Promotion of Science (Overseas Research Fellowship to TN) and the Swiss National Science Foundation (SNF: P2BSP2 168735 to JZ).

## 5. Appendix A

In the following, we show that our formulation of the astigmatic-defocus term as a quadratic form is equivalent to the traditional one as defined in RELION, which in turn was based on the model in CTFFIND (Mindell & Grigorieff, 2003). Let the two defoci be given by *Z*_1_ and *Z*_2_, the azimuthal angle of astigmatism by *ϕ*_*A*_ and the wavelength of the electron by λ. We then wish to show that:

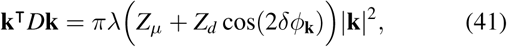

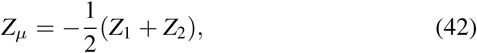

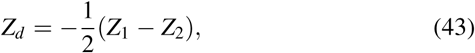

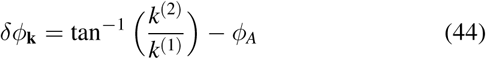

for the astigmatic-defocus matrix *D* defined as:

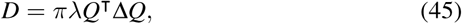

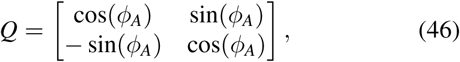

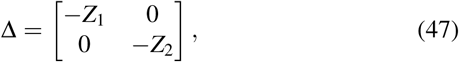

The multiplication by *Q* rotates **k** into the coordinate system of the astigmatism:

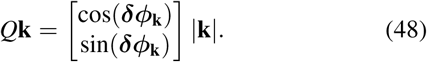

Multiplying out the quadratic form and applying the definitions of *Z*_*µ*_ and *Z*_*d*_ yields:

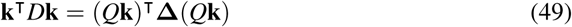

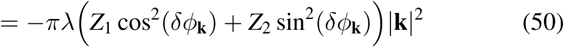

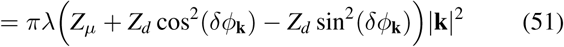

By substituting cos(2*δϕ*_**k**_) for cos^2^(*δϕ*_**k**_) − sin^2^(*δϕ*_**k**_) we see that this is equivalent to the original formulation.

In order to convert a given *D* into the traditional formulation, we perform an eigenvalue decomposition of − *D*/(*π*λ). The two eigenvalues are then equal to *Z*_1_ and *Z*_2_, respectively, while the azimuthal angle of the eigenvector corresponding to *Z*_1_ is equal to *ϕ*_*A*_.

## 6. Appendix B

Computing *T* and **l** explicitly through Eq. 38 would require iterating over all particles *p* in the data set. Since we have already accumulated the terms in *S*_***k***_ and **e**_***k***_ over all *p*, we can avoid this by instead performing the following summation over all pixels **k**:

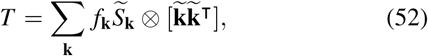

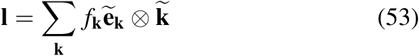

where ⊗ indicates element-wise multiplication, and the real symmetrical 4 × 4 matrix 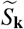 and the column vectors 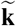 and 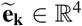 are given by the reshaping of *S*_**k**_, **k** and **e**_**k**_:

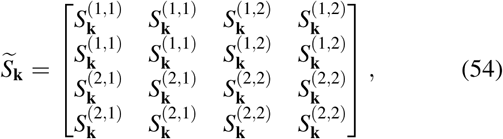

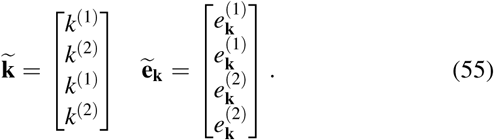

